# Entropy Transfer between Residue Pairs Shows that Allostery is an Intrinsic Property of Proteins: Quantifying Allosteric Communication in Ubiquitin

**DOI:** 10.1101/084756

**Authors:** Aysima Hacisuleyman, Burak Erman

**Affiliations:** Chemical and Biological Engineering, Koc University, Istanbul, Turkey

## Abstract

**Abstract:** It has recently been proposed by Gunasakaran et al. that allostery may be an intrinsic property of all proteins. Here, we apply Schreiber’s transfer entropy formulation to the non-allosteric protein Ubiquitin and show that there are indeed systematic pathways of entropy and information transfer between residues that correlate well with the activities of the protein. We use 600 nanosecond molecular dynamics trajectories for Ubiquitin and its complex with human polymerase iota and evaluate entropy transfer between all pairs of residues of Ubiquitin and quantify the binding susceptibility changes upon complex formation. Calculations show that specific residues act as entropy reservoirs in Ubiquitin and others as entropy sinks. Using the plausible conjecture that extracting entropy from a residue makes it more susceptible for interaction with a partner, we explain the ternary complex formation of Ubiquitin in terms of entropy transfer. Finally, we show that time delayed correlation of fluctuations of two interacting residues possesses an intrinsic causality that tells which residue controls the interaction and which one is controlled. Our work shows that time delayed correlations, entropy transfer and causality are the required new concepts for explaining allosteric communication in proteins.

**Author Summary:** Allosteric communication is essential for the function of proteins. Recent work shows that allostery results from dynamic processes in the protein associated with atomic fluctuations leading to entropic interactions that involve ensemble of pathways rather than discrete two state transitions. Based on this new picture of allostery, it was proposed that allostery may indeed be an intrinsic property of all proteins. In order to test this hypothesis, we derive the computational tools for quantifying allosteric communication, and explain allostery in terms of entropy transfer, a new concept based on information theory. We use long molecular dynamics simulations of proteins from which we calculate the transfer of entropy between pairs of residues. Results of simulations show that certain residues act as entropy sources while others as entropy sinks. Evaluation of time delayed correlations shows the presence of causality of interactions that allow us to differentiate between residues that are drivers in allosteric activity and those that are driven. Identification of driver-driven relations is important for drug design. Using the example of Ubiquitin, a protein that is not known to be allosteric, we identify paths of information transfer that control its binding to diverse partners in the Ubiquitin-Proteasome System. We conclude that allosteric communication resulting from entropy transfer between residues is an intrinsic property of all proteins.

## Introduction

Allosteric communication describes the process in which action at one site of a protein is transmitted to another site at which the protein performs its activity. The importance of allostery in biological systems has generated significant experimental and computational research. The basic problem is to identify residues that participate in allosteric communication in the hope of controlling their behavior related to protein function. Allosteric communication first requires the identification of two sites, the effector site, i.e., the site that is acted upon, and the regulatory site where protein’s activity is regulated. Although more than 1000 allosteric sites are known [1] many more need to be characterized. In fact several pairs of allosteric endpoints may exist in a protein [2] which increases the number of candidate pairs that communicate. This problem becomes even more important when one considers the fact that most known cancers result from disruption of allosteric communication as a result of single mutations [3, 4] and the number of proteins associated with this phenomenon is very large. Expressed in simple terms, the solution of the problem reduces to finding whether two given residues communicate with each other, and if so what the consequences of this communication are. Various approaches to solve the problem may be found in References [5-16]. The specific aim of the present paper is to develop a rapid computational technique that identifies interaction of residue pairs based on concepts of information transfer and entropy, to scan a given protein and identify pairs of sites that communicate and to determine whether these communicating pairs may be candidates for allosteric activity.

The present work departs from the approaches outlined in the preceding paragraph. We do not focus neither on single allosteric sites nor on allosteric paths. We consider the time trajectory of the fluctuations of two residues, which may be spatially distant, and search for information transfer from the trajectory of one residue to that of the other. The trajectories are obtained from long molecular dynamics (MD) equilibrium simulations that give the fluctuation of each atom at constant temperature. The first requirement for information to be transferred from an atom *i* to another atom, *j*, is that their trajectories should be correlated. The second requirement is that this transfer should be asymmetric, i.e., information going from *i* to *j* should not be equal to information from *j* to *i*. This requires the use of time delayed correlations of fluctuations which may be asymmetric in contrast to time independent correlations which are symmetric by definition and therefore lack information on directionality. If *C_ij_*(t,t+τ) denotes the correlation of fluctuations of *i* at time *t* with those of *j* at time *t*+τ, then asymmetry requires that *C_ij_*(t,t+τ)≠*C_ij_*(t,t+τ). This introduces directionality and therefore causality into the problem. If time delayed correlations are asymmetric, then can we quantify the net information that is transferred? The answer is yes if we pose the problem in terms of entropy transfer.

Before going into the discussion of entropy, it is worth pointing out that information transfer is exclusively based on the changes in the amplitudes and frequencies of fluctuations in the system. This was first suggested and modelled by Cooper and Dryden (CD) [17] and reached larger dimensions by the work of Gunasekaran et. al. [18] which suggests that since allosteric communication is a result of correlated fluctuation then allostery should be an intrinsic dynamic property of all proteins. The dynamics aspect of proteins resides in the fluctuations of atoms which may be evaluated by experimentally measuring the B-factors of the atoms. The B-factor of the ith atom is related to its time independent autocorrelation of fluctuations, 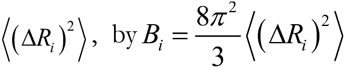. However, knowledge of them is not sufficient for predicting allosteric communication and cross correlations 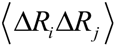 between the fluctuations of different atoms are needed. Allosteric activity requires not only the modulation of the cross correlations in the system but also on time delayed cross correlations,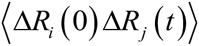 as will be described below in detail. The CD model is referred to as ‘Allostery without conformational change’. In this respect, it goes beyond the classical Monod-Wyman-Changeux (MWC) [19] model and its relative, the Koshland Nemethy Wyman (KNW) model [20] both of which relate allostery to discrete conformational changes at the regulatory site. Sending information by changing the amplitude and frequencies of fluctuations is entropic [21] and depends not only on the value of the entropy but also on the transfer of entropy from residue to residue during communication. Entropy as a source of information transfer is widely used in information theory [22] which is only very recently used for a protein-DNA complex by Kamberaj and van der Vaart [23]. Through analysis of entropy transfer, they determined residues that act as drivers of the fluctuations of other residues, thereby determining causality that is inherent in the correlations. Determining residues that act as drivers and those that are driven is important especially from the point of view of drug design. Entropy transfer and causality is a new paradigm for studying allosteric communication in proteins, which we elaborate in detail in the present paper. On a broader scale, our findings show that all proteins may indeed exhibit allosteric communication and therefore supports the hypothesis by Gunasakaran et. al., [18] which states that allostery is an intrinsic property of all dynamic proteins.

The quantitative measure of information flow between two correlated processes is introduced by Schreiber [22] in 2000. In the present work, the processes are generated in the form of trajectories of atomic coordinates using MD simulations from which probabilities of atomic coordinate fluctuations required for evaluating transfer entropy are calculated. We calculate the entropies based on atoms and identify the entropy of a residue with the entropy of its alpha carbon. Denoting the probability of fluctuation of atom *i* by *pi*, Callen showed [24] that the Shannon measure of disorder, 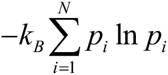 with *N* and *k_B_* denoting the number of elements of the system and the Boltzmann constant, is the entropy of the system which is maximized at constant energy (See Callen [24], Chapter 17. Entropy and disorder: Generalized canonical formulations). At this point we give here a qualitative explanation of the relationship between information flow and a physical event such as fluctuations of atoms, and continue this discussion on a quantitative basis after we introduce the statistical mechanical basis of the model. Suppose we have two trajectories, one of atom *i* and the other of atom *j*. If the fluctuations of *i* and *j* are independent of each other, than knowledge of the fluctuations of i will not give us information on the fluctuations of *j* and the uncertainty associated with the two events will be a maximum. The total entropy of i and j will be the sum of the singlet entropies, *S*_*i*_ + *S_j_*. If, on the other hand, i and j move in a correlated way, the fluctuations of *i* controlling the fluctuations of *j*, then we will have more information on the fluctuations of *j* than if they were uncorrelated. For example, if *i* and *j* were perfectly correlated, then we would know exactly what j will do if we know what i is doing. This extra information *I_ij_* that we gain because of the physical coupling of *i* and *j* is obtained by the Shannon equation and is termed as the mutual information and is always positive. The total entropy, *S_ij_* of *i* and *j* in this case is written as *S_ij_* = *S_i_ + S_j_ - I_ij_* (see Eqs 12 and 13 and also Ref. [25]). Thus, correlation of fluctuations, irrespective of whether they are negative or positive, always decreases the sum of the singlet entropies of *i* and *j*. These arguments and the Shannon equation have been used to obtain entropy changes in proteins at different levels of approximation [21, 26-38]. However, we need to go beyond the Shannon equation in order to quantify allosteric communication in proteins which requires, as shown by Schreiber in 2000 [22], the knowledge of time delayed conditional probabilities of two trajectories. In the interest of determining which residue drives the correlated motions and which residue responds, van der Vaart applied the Schreiber equation to determine information flow between Ets-1 transcription factor and its binding partner DNA [23] (Also see references [39] and [40] in similar context). Since this first work on entropy transfer in proteins there has been a limited number of studies on information transfer in proteins. Barr et al. [41] quantified entropy transfer among several residues in a molecular dynamics analysis of mutation effects on autophosphorylation of ERK2. Corrada et al. [42] analyzed entropy transfer in antibody antigen interactions. Perilla et al. [43] used the transfer entropy method to analyze barrier crossing transitions in epidermal growth factor receptors. Qi and Im [44] quantified drive-response relations between residues during folding. Jo et al. [45] obtained a causality relationship between intramolecular hydrogen bonds and the conformational states of *N*-glycan core in glycoproteins. Zhang et al. [46] applied the method to understand changes in the correlated motions in the Rho GTPase binding domain during dimerization. An extensive overview of similar techniques is given in reference [47].

In the following section, we define the model on which we build the information theoretical basis of entropy. We then study the problem of time delayed correlation of fluctuations in proteins. Despite its importance in pointing to directionality of events in proteins, as has been shown recently for the allosteric activity od K-Ras [48], time delayed functions have not been studied in detail in the past. We then present a fast and accurate method of calculating entropy changes in proteins subject to pairwise interactions. Calculation of entropy of proteins is not new and has already been investigated by several authors [26-28, 49, 50] at different levels of approximation. Our method of entropy calculation is motivated by the recent finding that the distribution functions for the magnitude of fluctuations of residues in globular proteins can be derived from a universal function [51]. The method that we use for calculating the entropy is fast and accurate, based on histogramming the magnitude of fluctuations of each atom in a protein where the bin number is chosen according to the Sturges’ rule of determining the widths of class intervals [52]. We show that the use of Sturges’ rule in our computational method leads to results that are in agreement with earlier entropy calculations. We benchmark our method with calculations of Ubiquitin by Fleck et al [38]. The entropy change of Ubiquitin upon binding to human polymerase iota UBM2 that we calculate gives the same value obtained in reference [38] using a different method of entropy estimation. The computational method that we adopt is efficient and plausible.

The association of Shannon equation with statistical mechanical definition of entropy and quantifying transfer entropy by the use of the Schreiber equation allows us to interpret a wide range of events in proteins. If entropy transfer is considered in terms of changes in mobility, then transfer of entropy from *i*to*j*implies decrease in the mobility of *i* due to its correlation with *j*. Stated in anotherway, residue*j*extracts entropy from *i*. If binding is considered, one could then say that transfer ofentropy from*i*to*j*would facilitate binding at *i* due to lowered mobility of *i*, although this may not be a general trend and may depend on several other factors. We use the model to study the directionality of information flow and entropy transfer in the 76 amino acid protein Ubiquitin which is known to propagate signals allosterically in the cell by binding to a vast number of substrates [53]. Ubiquitin itself is not known to be allosteric, however, its coordinated motions upon binding with a protein modifies the fluctuation patterns on another site that affects the binding of a third protein as in the case of its ternary complex [54]. In order to identify communication patterns leading to such effects, we scan the full Ubiquitin and identify the pairs of residues whose time delayed correlation functions are asymmetric and quantify the amount of entropy transferred between these residues. We then analyze the behavior of Ubiquitin when complexed with the binding partner human polymerase iota UBM2, 2KTF.pdb.

## Results

### Structure of Ubiquitin

Ubiquitin is a 76 amino acid protein as shown in Fig 1. It consists of 8 distinct secondary structures that actively take part in its interactions with a large number of proteins.

**Fig 1.**
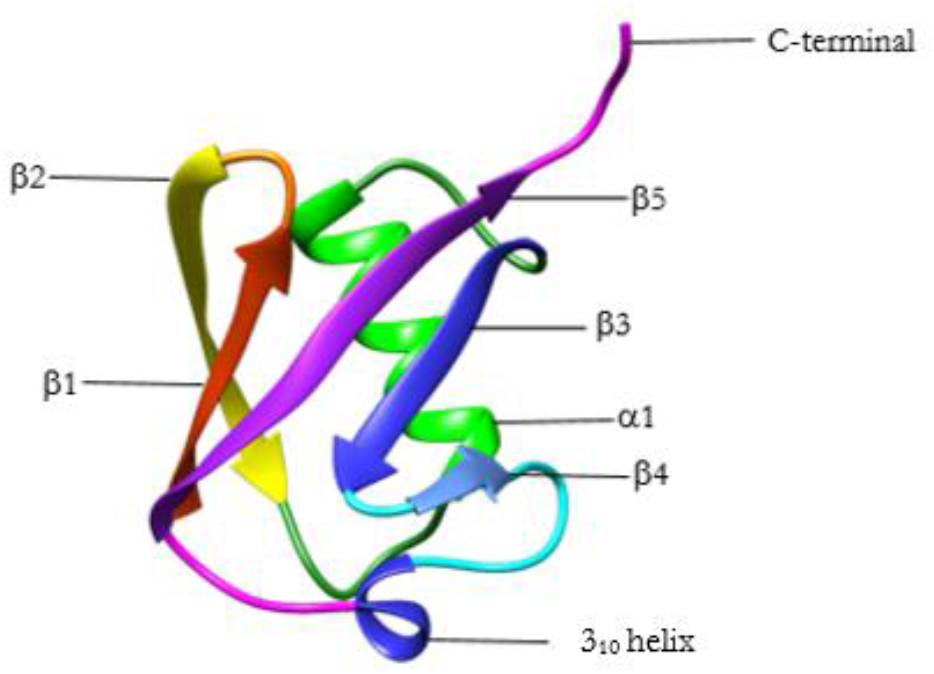
*Structure of Ubiquitin. Ribbon diagram was generated by Chimera.*

The interactions of the secondary structures are strictly coordinated by the correlations in the protein. In Fig 2 we present the results of Pearson correlations of fluctuations, where the negative and positive correlations are shown in the left and right panels, respectively. The correlations with amplitudes (-1,-0.25) and (0.25, 1.0) are shown in the figure. The negative correlation map shows that β1 is correlated with α1, β3 and β4. It is also correlated with the loops between β1- β2, 1-β3 and β3-β4. β2 is correlated with β4 and C-terminal. It is also correlated with the loop between β2-α1 and β4-β5. β1 is correlated with β1, β2, the loop between and 1 and the C-terminal. β3 is correlated with β1, β2, the loop between β4 and β5 and the C-terminal. β4 is correlated with β1, β2, C-terminal and the loop between β4-β5. β5 is correlated with the loop between β4-β5 and the C-terminal. The positive correlation map shows that β1 is positively correlated with β2, β5 and C-terminal. It is also correlated with the loop between β4 and β5. β2 is positively correlated with β1. α1 is positively correlated with β3, β4, β5 and the loop between α1- β3, β3-β4 and β4- β5.β3 is positively correlated with α1, β4 and β5. α1 is also positively correlated with the loop between β1-β2, α1-β3 and β3- β4. β4 is positively correlated with α1 and β5. β5 is positively correlated with β1, α1, β3, β4 and the loop between β1-β2.

**Fig. 2.**
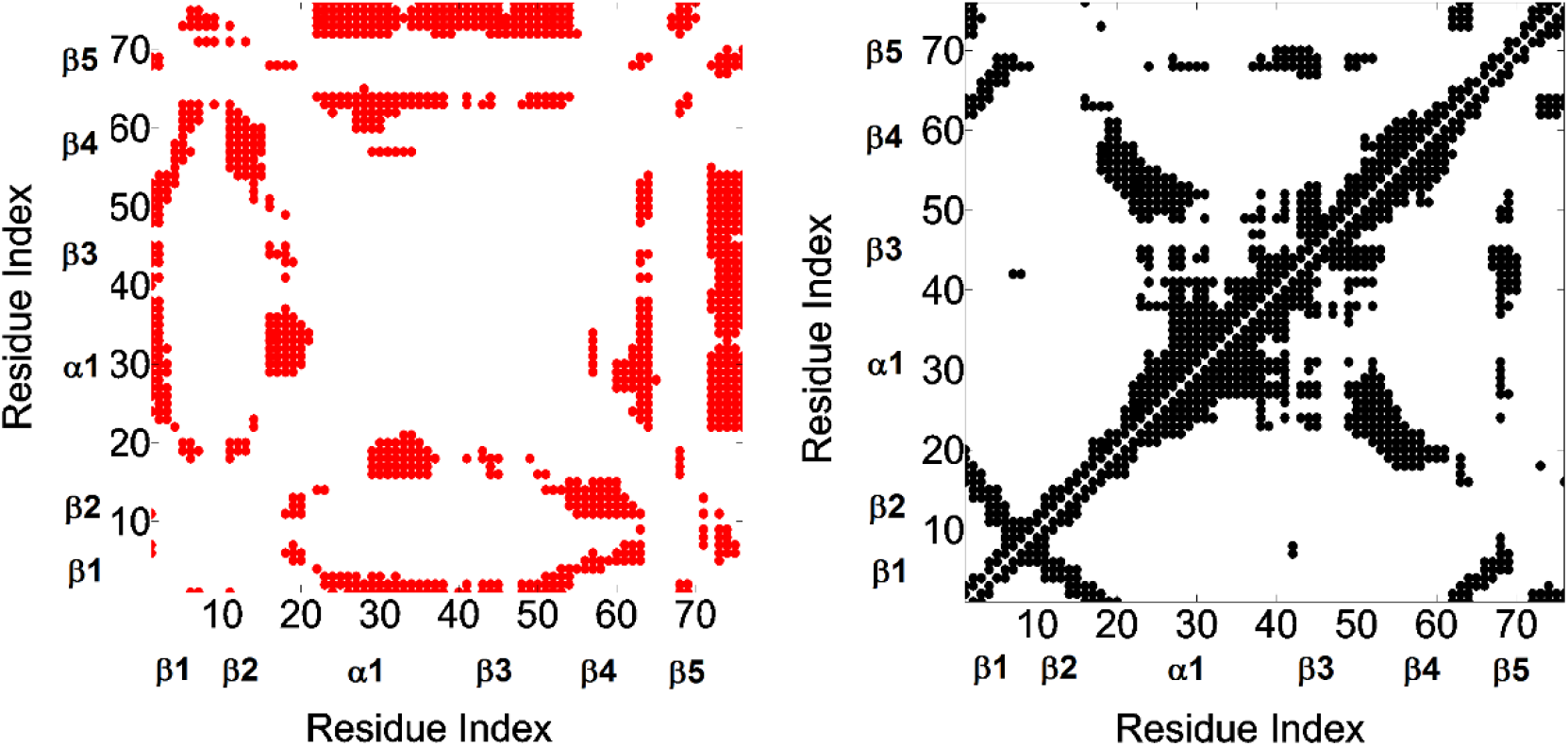
The left panel shows the negative Pearson correlations in the range (−1, −0.25), the right panel shows positive Pearson correlations in the range (0.25, 1.0). Pearson correlations are calculated from Eq. 9.

### Transfer entropy in Ubiquitin

We present the results of entropy transfer between all residue pairs of Ubiquitin. We consider only the alpha carbons, and the values given are divided by the Boltzmann constant. Using Equation 16 we evaluated the values of entropy transfer from alpha carbon i to j, *T_i→j_*(τ), for all pairs of i and j for τ= 5 ns. Calculations averaged over several time stations between 0 and 5 ns gave approximately the same values for entropy transfer. In the remaining parts we present τ= 5 ns results only. The characteristic decay time of correlations of fluctuations of alpha carbons, which will be discussed in the following section, is on the average between 5 to 10 ns. The entropy transfer function *T_i→j_*(τ) that we obtain from fluctuation trajectories of alpha carbons depends on the correlation of two events that are τ ns apart in time. If τ is taken very small, i. e., around zero, then the difference between *T_i→j_*(τ) and *T_j→i_*(τ) will be very small because *T_i→j_*(0) = *T_j→i_*(τ). If is taken much larger than the characteristic decay time, then the correlations will have decayed to small values and the differences will be vanishingly small. In agreement with this reasoning, we took τ=5 ns and calculated entropy transfer at this time. The results are shown in Fig 3. The abscissa, named as entropy donor, denotes the indices of residues that act like entropy reservoirs to other residues. The ordinate, named as entropy acceptor, denotes the indices of residues that act like entropy sinks that absorb entropy from the system.

**Fig 3.**
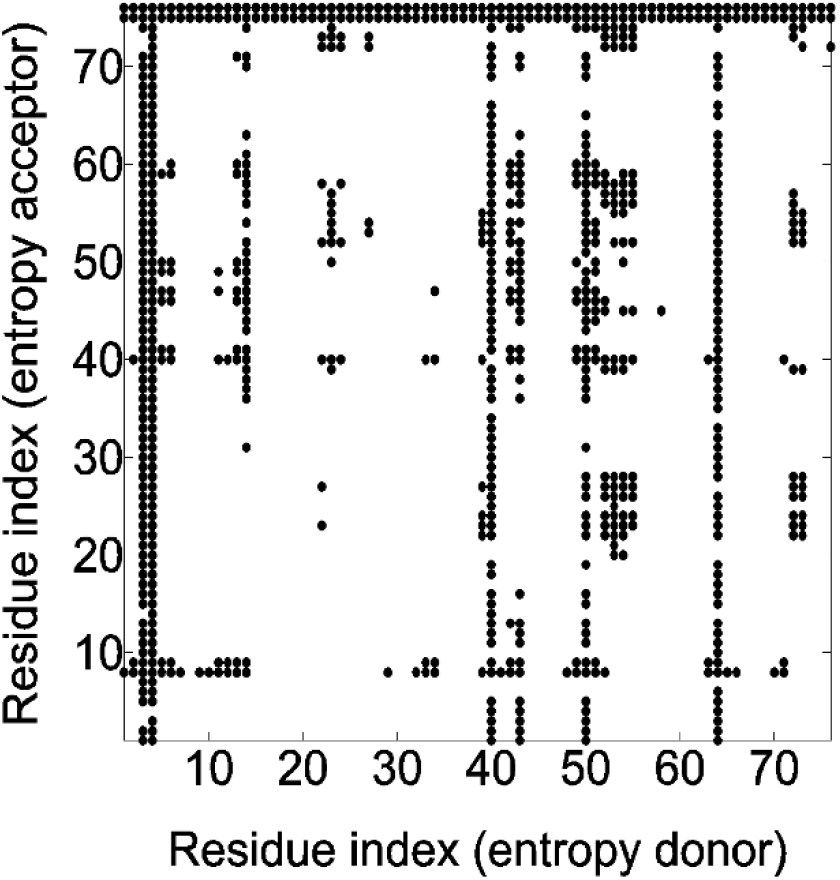
*Entropy transfer from residue i to residue j. Abscissa represents residues which provide entropy to residues shown along the ordinate. Entropy transferred from residue i to residue j is obtained from Eq. 16. Values between 0.0035-0.35 are recorded. Values below 0.0035 are not shown in order not to crowd the figure. T_i→j_*(τ) values are calculated from Eq. 16 with τ=0.5 ns.

The columns of black points in the figure show that specific residues, such as ILE3 and PHE4, ILE13, ILE23, LYS27, GLY53, GLU64, ARG72 provide entropy to several residues of the protein. The rows of black circles indicate residues such as LEU8, THR9, GLY75 and 76, that absorb entropy from several residues of the protein. Residues ILE3 and PHE4, ILE13 and GLU64 form a spatial cluster. Also, the residues ILE23, LYS27 and GLY53 form a spatial cluster. If the allosteric path description is adopted, then we can say that these two spatial clusters lie on the allosteric path.

The net transfer of entropy from residue i, defined by Eq. 17 is presented in Fig 4. Positive values denote net entropy transfer out from a residue, and negative values denote net entropy into a residue. Similar to the pattern observed in Fig 3, we see that certain residues behave as entropy sources for the rest of the protein and some behave as entropy sinks.

**Fig 4.**
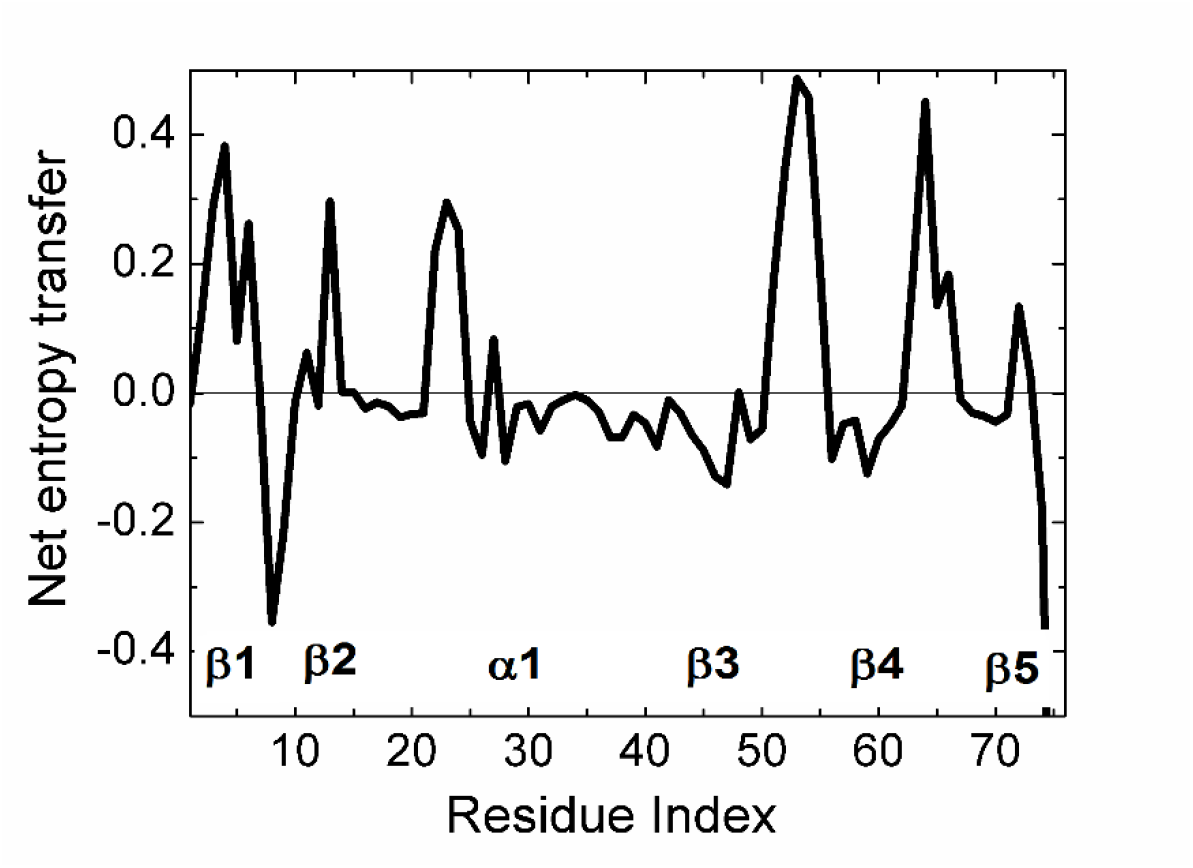
*Net entropy transfer from one residue to the rest of the protein, calculated by Eq. 17.*

We see that β1 and β2 act as an entropy sources as well as part of the helix α1. The largest amount of entropy is provided by the loops between β3 β4 and β4 β5. The two major entropy sinks are the loop between β1 and β2 and the last two residues of the C-terminal. Entropy sources are located mostly at secondary structures or at their extremities. The three residues PHE4, THR14, GLU64 are spatial neighbors. Similarly, LEU43, LEU50, are spatial neighbors.

In order to have an idea on the mechanism of communication in the system, one needs to know the transfer of entropy among specific pairs of residues. From the data of Fig 4, we can find with which residues a given amino acid interacts entropywise. Fig 5 and 6 summarizes the net entropy exchange, *T_i→j_*(5)–*T_j→i_*(5), between the labeled residue in each panel and the j’th residue of the protein. Fig 5 shows some examples with mostly positive entropy transfer from the labeled residue to others. The top left panel in Fig 5 shows entropy transfer from ILE3 to other residues. Specifically it transfers the largest entropy to LEU8 and GLY75 and GLY76. From the dynamics point of view, we can say that ILE3 contributes to the mobility of these three residues. Both ILE3 and LEU8 are at the opposite extremities of β_1_. ILE3 is a spatial neighbor of GLU64. GLU64 is hydrogen bonded to GLN2 which is on a loop, and the mobility of the loop is transferred to ILE3 via the stated hydrogen bond. ILE3 also contributes entropy to several other residues of the protein as may be seen from the figure. Entropic interactions of residues PHE4, ILE13, ILE23 and LYS27 are very similar to those of ILE3 and are not shown in Fig 5. The top right panel in Fig 5 shows the interactions of GLY53 with the rest of the protein. GLY53 is situated on the long loop between β_3_ and β_4_, and is hydrogen bonded to the main alpha carbon of GLU24 which is at the end of α_1_. Fig 5 shows that GLY53 contributes to the mobility of the segment between VAL17 and LYS29. It also transfers entropy along the chain to LEU56. GLU64 contributes entropy to several residues, in a way similar to that of ILE3. ARG72 has a unique pattern of contribution, specifically to ASP39 which is its spatial neighbor, to the loop between α_1_ and β_2_, to PHE45 and LEU56, both of which are spatially distant from ARG72. It also contributes to the mobility of the C-terminal. Fig 6 gives two examples for mostly negative values of *T_i→j_*(5)–*T_j→i_*(5). The left panel in Fig 6 shows that LEU8 and GLY76 absorb entropy from most of the residues of the protein.

**Fig 5.**
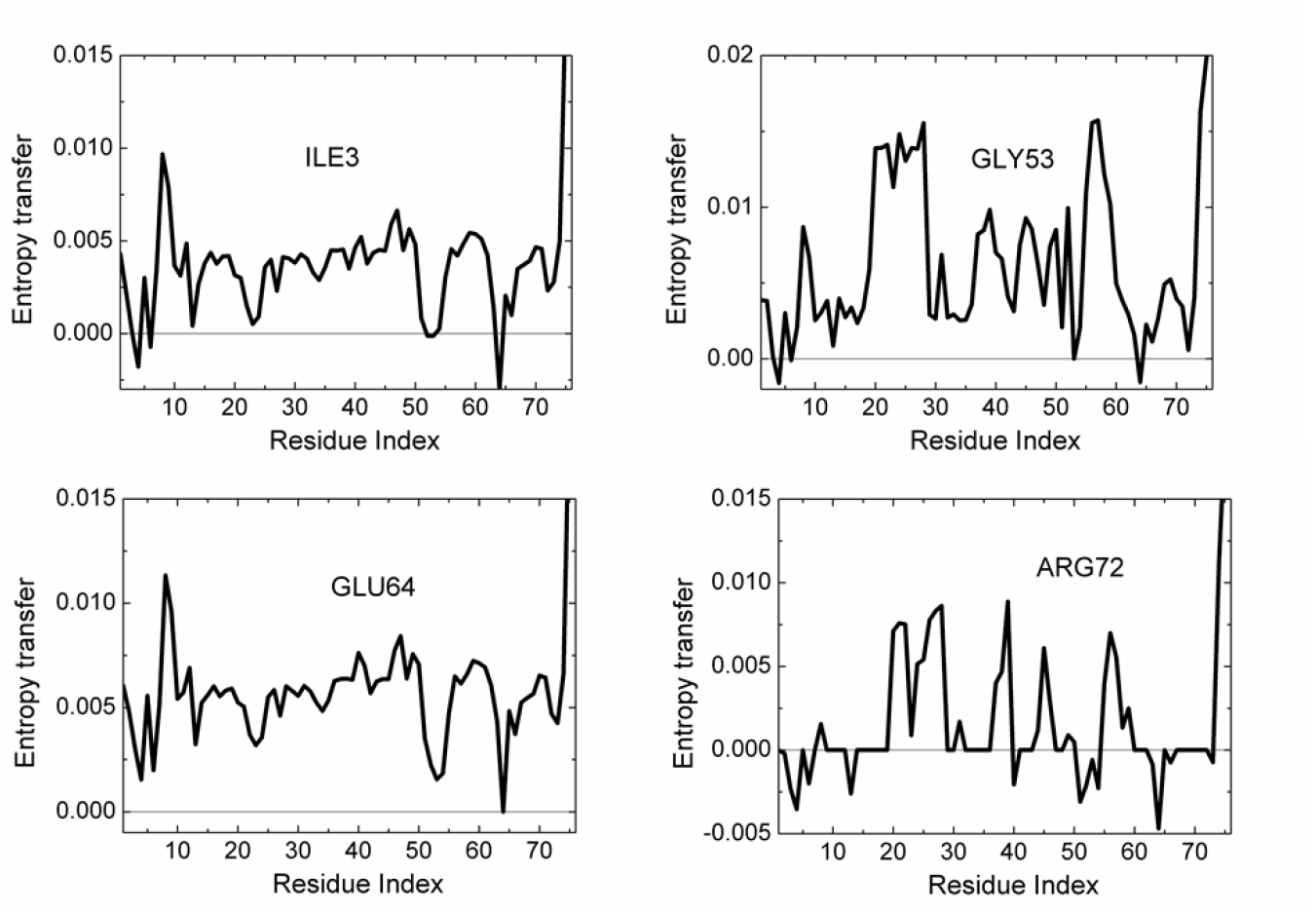
*Entropy transfer from a given residue to other residues of the protein. The residue from which entropy is transferred is marked in each panel. Calculations are based on the relation T_i→j_*(5)–*T_j→i_*(5).

**Fig 6.**
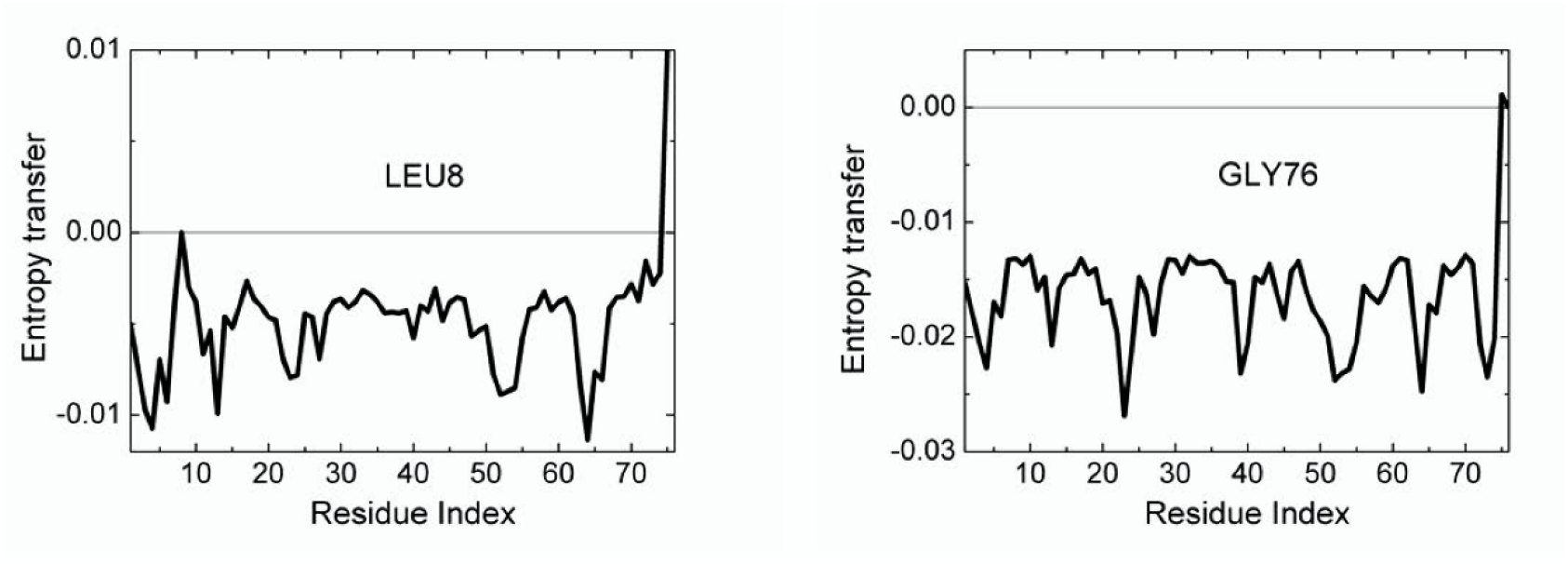
*Entropy transfer from residues of the protein into LEU8 and GLY76*. *Calculations are based on the relation T_i→j_*(5)–*T_j→i_*(5).

### Time delayed correlations of Ubiquitin

Fluctuations of amino acids in Ubiquitin display characteristic decay times that are in the order of 1 to 5 ns. Differences arise from the unique conformational features of the amino acid and its environment. In Fig 7, we show the autocorrelations of LEU7 and THR71.

**Fig 7.**
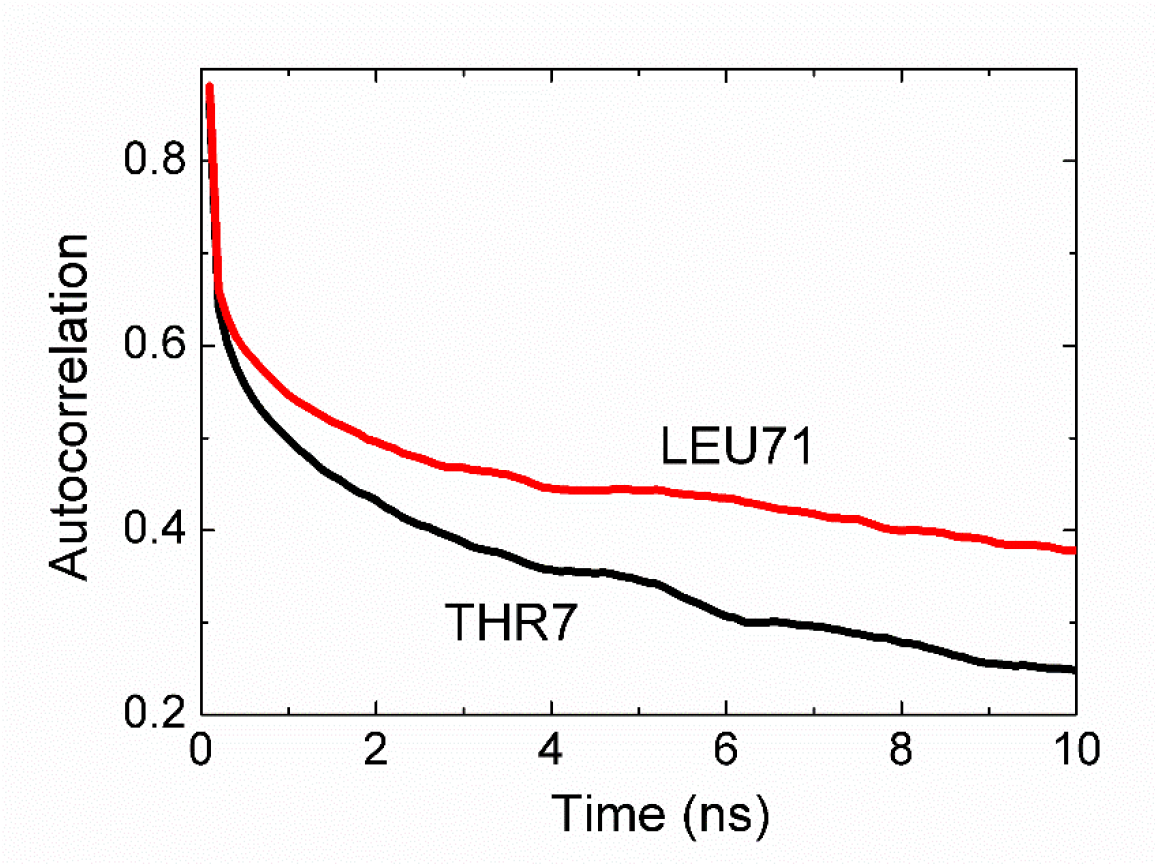
*THR7-LEU71 Autocorrelations calculated from Eq. 7 for i=j.*

The autocorrelation function for THR7, i.e., the time required to decay to 1/e of the original value is 5 ns. LEU71 decays slightly slower with a decay time of 10 ns¨

The time delayed cross correlations of the fluctuations of two amino acids are of interest because yield information on the causality of events. The static correlations presented in Fig 2 are symmetric, *C_ij_*(τ)=*C_ji_*(τ). However, time delayed cross correlations of fluctuations of two amino acids show asymmetries which we discuss in this section.

The strongest asymmetry is between LEU7 and THR71, shown in Fig 8.

**Fig 8.**
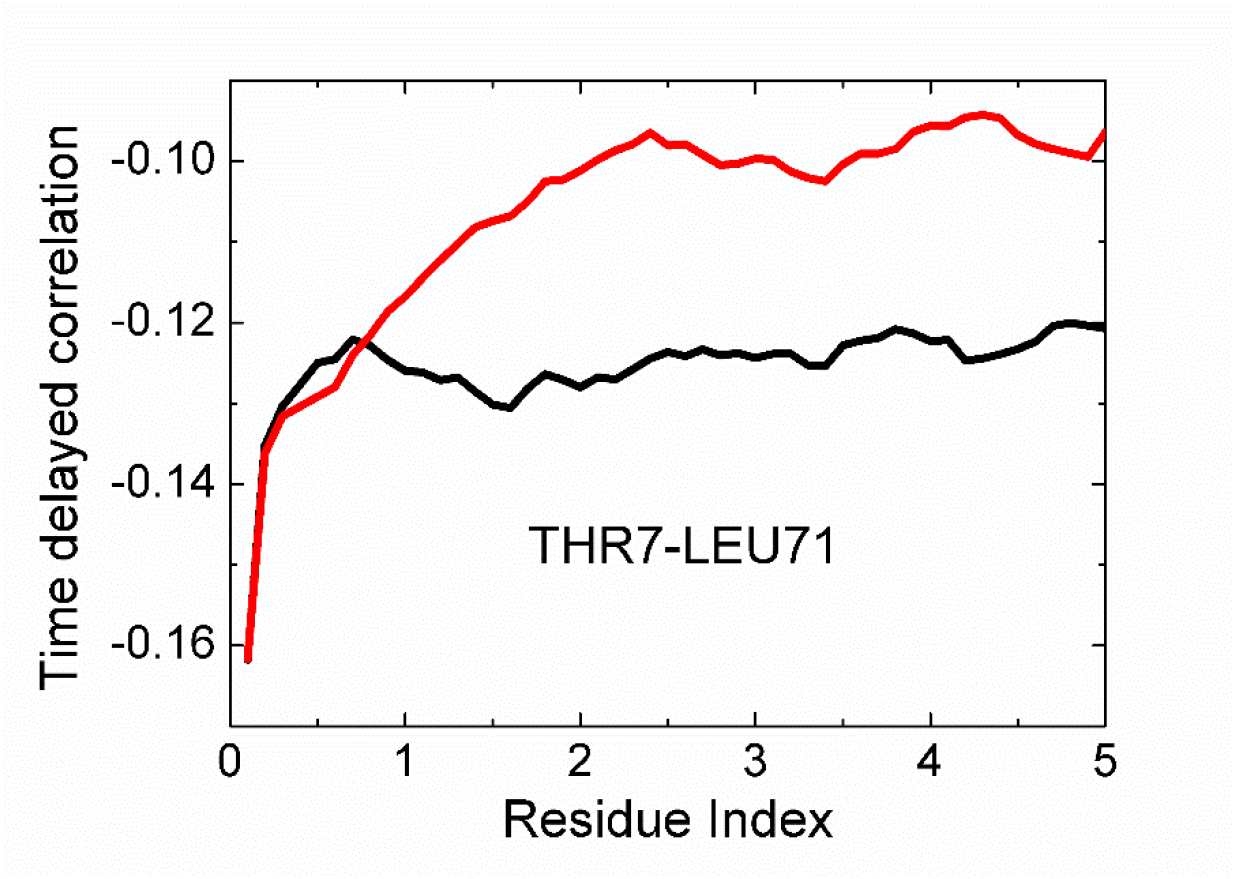
*Cross correlation of fluctuations of THR7 and LEU71. Black line is for correlations where THR7 precedes LEU71. The red line is for correlations where LEU71 precedes THR7. The curves are calculated from Eq. 7.*

In this figure, the black curve is for correlation of THR7 at time t and LEU71 at t+τ. The red curve is for LEU71 at t and THR7 at t+τ. The black curve decays significantly slower than the red curve, indicating that the effect of the fluctuations of THR7 on later fluctuations of LEU71 persists for longer times whereas the converse is not true. We therefore say that the motions of THR7 drive the motions of, LEU71 i.e., THR7 is the driver and is LEU71 driven. Since LEU71 is located on the C-terminal segment, and THR7 is at the end of 1, we can say that the 1 strand controls the fluctuations of the C-terminal. We see that the black curve remains approximately constant after a rapid initial decay. This shows that the driver action of THR7 on LEU71 persists for longer times.

**Fig 9.**
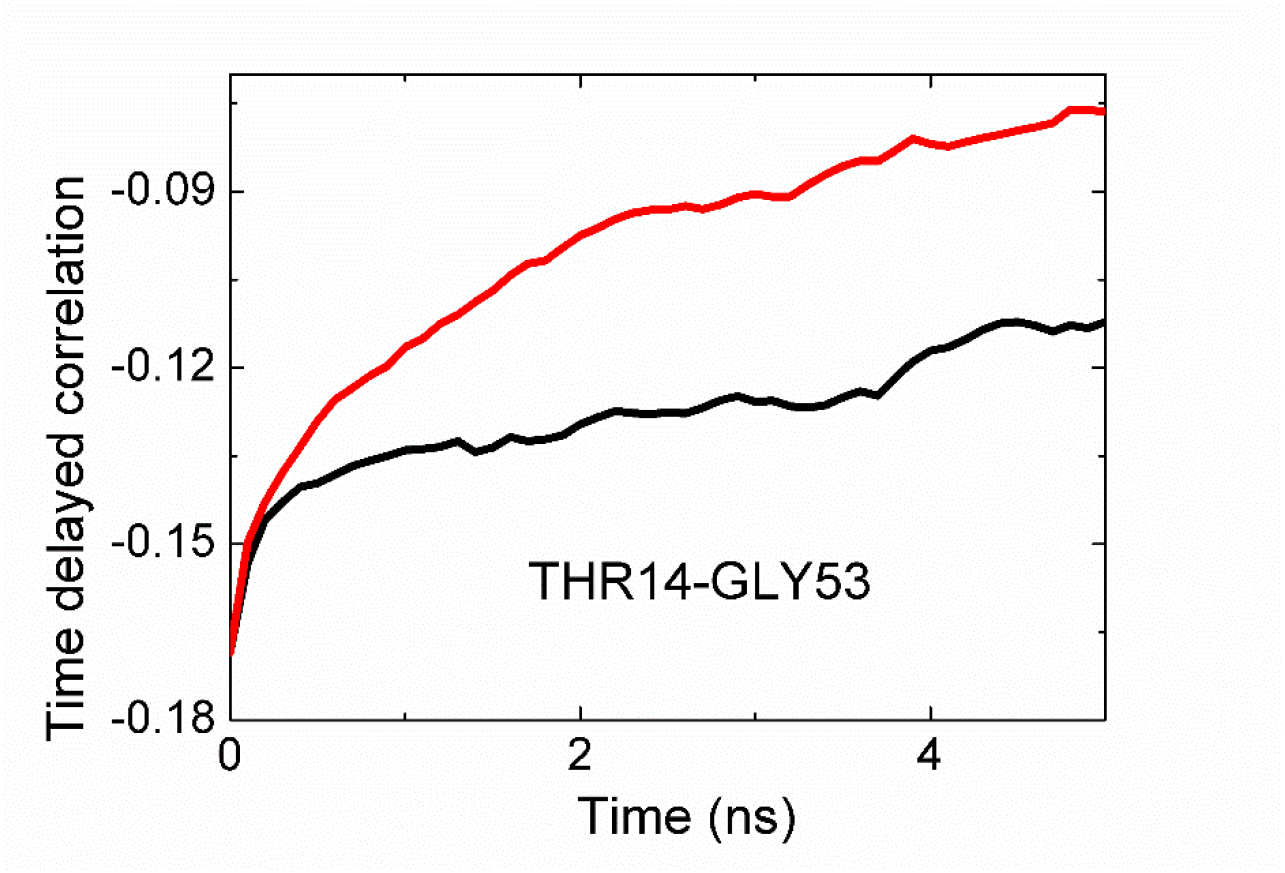
*Cross correlation of fluctuations of THR14 and GLY53. See legend for Figure 8.*

In Fig 9, the black curve is for the correlation of the fluctuations of THR14 with later fluctuations of GLY53. The red curve is for the reverse case, fluctuations of GLY53 affecting later fluctuations of THR14. This figure shows that THR14 is the driver and GLY53 is driven. THR14 is on the 2 strand and GLY53 is on the long loop connecting the 310 helix to β4.

### Changes upon complex formation of Ubiquitin

Ubiquitin forms complexes with a multitude of proteins. Here we studied its complex with Human Polymerase Iota (UBM) which is a small protein of 28 amino acids, 2L0G.pdb. UBM binds to Ubiquitin at β_1_, the loop between β1 and β_2_, and at β_3_ and β_4_. The binding affects the fluctuations of these regions. We compare the B-factors for Ubiquitin in the bound and unbound states in Fig 10. The solid and the dashed curves for Ubiquitin in the bound and free states, respectively. Binding decreases the B-factors of the contact substructures as may be seen from the figure. Additionally, the fluctuations of certain residues between 23 and 37 are decreased although UBM binds to a region far from these residues. Interestingly, the amplitude of fluctuations of the C-terminal residues are not affected by the binding.

**Fig 10.**
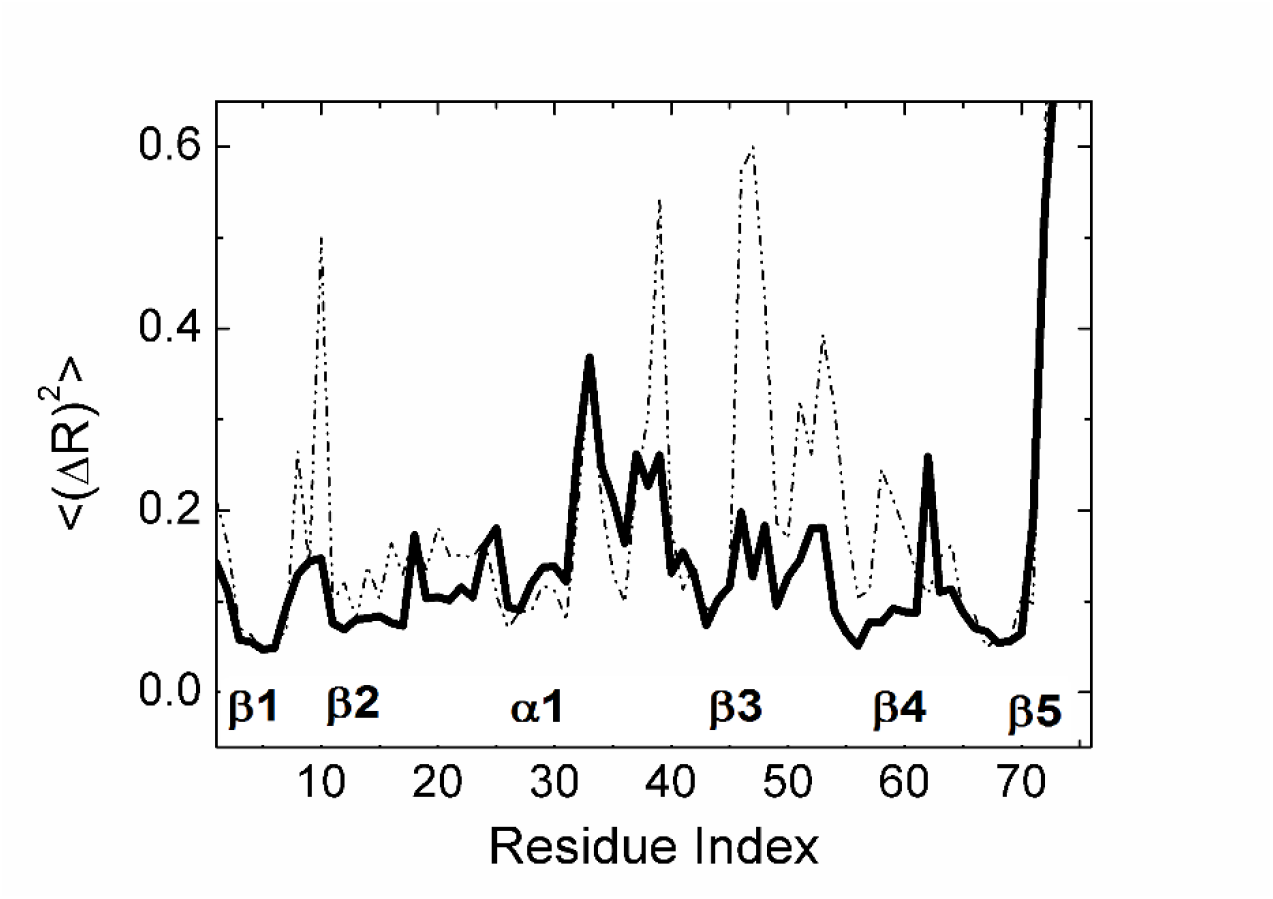
*Mean-square fluctuations of residues in unbound Ubiquitin (dashed curve) and in the complex (solid curve)*

Net entropy transfer in Ubiquitin in the bound and free states is compared in Fig 11. The solid and dashed curves are for the bound and free states, respectively. We see from the two curves that Ubiquitin upon complex formation becomes a totally different structure from entropy point of view. Entropy transfer associated with residues ILE3, PHE4, ILE23, GLY53 and GLU64 is diminished and entropy transfer from LEU8 is markedly increased. ILE13 is not affected. Entropy transfer associated with the last two residues of the C-terminal is strongly decreased although their B-factors are not affected by binding.

**Fig 11.**
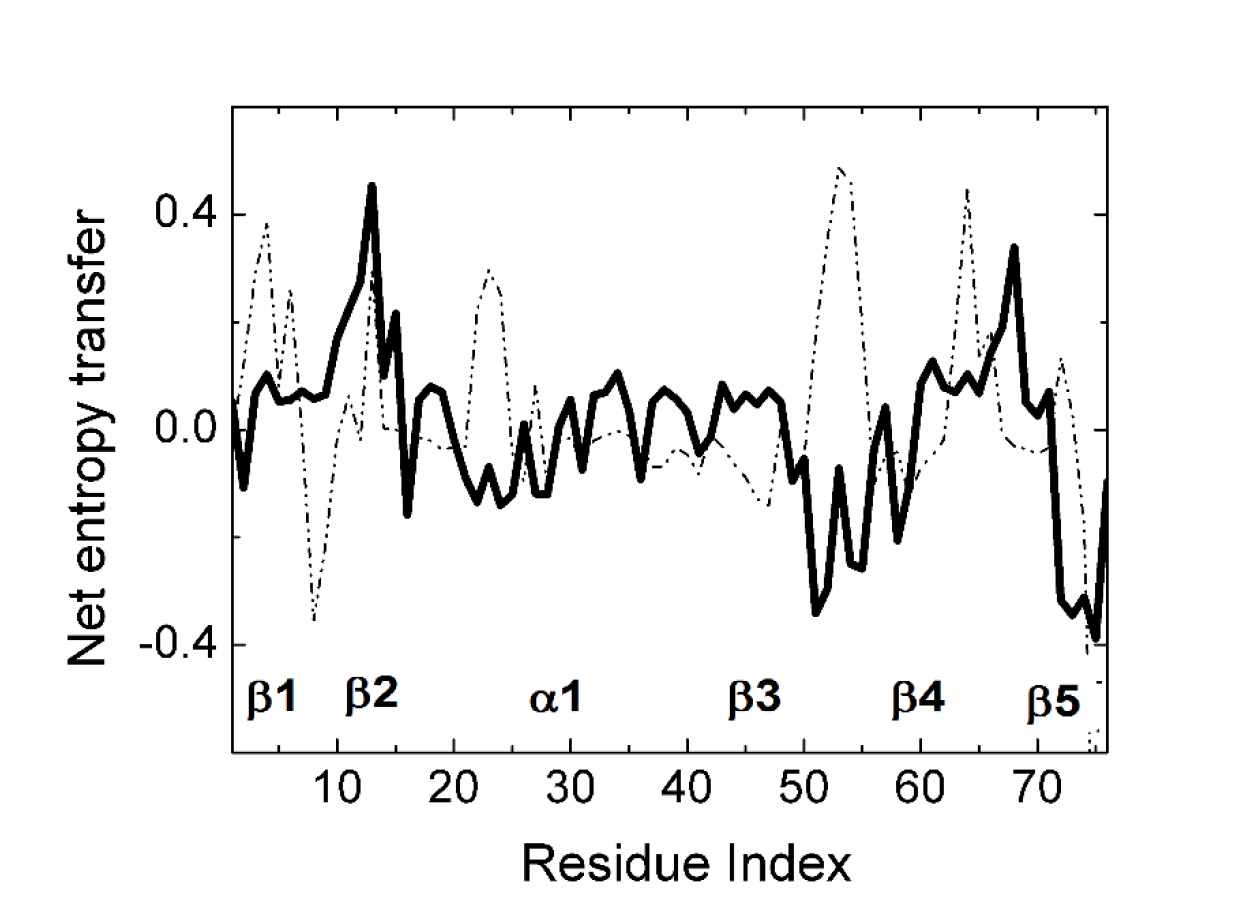
*Net entropy transfer of residues in unbound Ubiquitin (dashed curve) and in the complex (solid curve) Curves are obtained by using Eq. 17.*

Residues on the two loops between β_2_ and α_1_ (from SER20 to ASN25) and between β_3_ and β_4_, (from LEU50 to ASN60) are not involved in binding with UBM. These two regions, which were entropy sources in the unbound state now become entropy sinks, as may be seen from Fig 11 where the dashed curves have positive values and the solid curves have negative values in these two regions. Based on the arguments presented in this paper, we can say that these regions with entropy sink features will facilitate binding.

## Methods

### Molecular dynamics trajectories

We perform molecular dynamics simulations for a protein in equilibrium and extract stationary trajectories for each atom. The trajectories for the atoms are expressed as

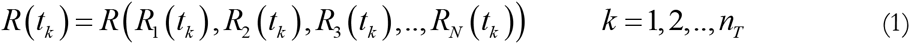

Here, *R_i_*(*t_k_*) is the position vector of the ith atom at the kth time tk, expressed in terms of its Cartesian coordinates, *X*_*i*_ (*t_k_*), *Y_i_* (*t_k_*) and *Z*_*i*_ (*t_k_*), *N* is the total number of atoms and *t_k_* is the time in the kth step. *k* ranges from 1 to *n_T_*, the total number of steps in the simulation. If the total time is *T*, then the *T*, length ξ of each time step is ξ=*T* / *n_T_*. Each atom has a unique equilibrium mean position defined by

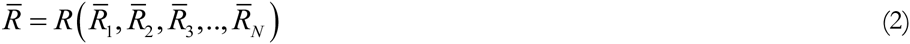

We denote the instantaneous state of fluctuation of a protein at time *t_k_* by the vector

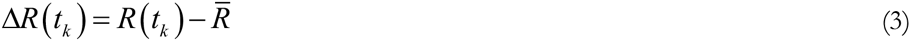

which reads in vector form as

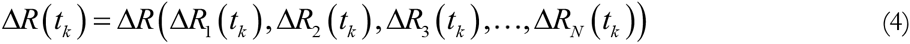

For each*t_k_*, Eq.4 has *N* entries. For the purposes of the present study, we need the magnitude of the fluctuations only. In the following, we will let *ΔR_i_*(*t_k_*) represent the magnitude of the fluctuation at time *tk*.

### Evaluation of probabilities

The most general expression for the probability of fluctuation Δ*R* is the joint probability *p*(*ΔR*) *p*(*ΔR_1_,ΔR_2_,ΔR_3_,…,ΔR_N_*). This expression contains information on all orders of dependence between atoms and is too general for use. In the other extreme, the simplest expression is the singlet probability function *pi*(Δ*R*_i_) which is obtained from the most general expression by

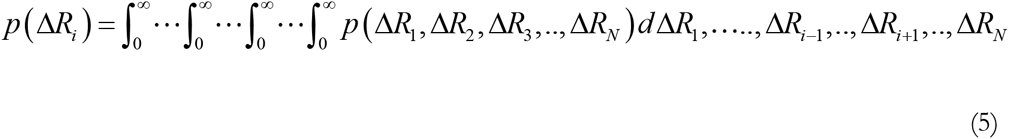

*N* such functions define the probability of fluctuations of the *N* residues within the singlet approximation.

The next simplest probability is the pair probability *p(ΔR_i_*,*ΔR_j_)* obtained from the most general expression by

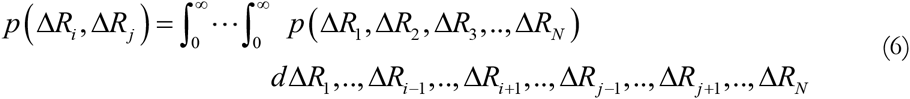

For *N* atoms, there are 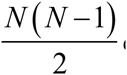 equations for pair probabilities.

In Eqs. 5 and 6,*ΔR*’ *s* are treated as continuous. In the remaining of the paper, we will adopt a discrete representation for them in terms of histograms. The histograms will be expressed in terms of *n* bins. We refer to each bin as a state. The variables in the probabilities will then be functions of state variables. Thus we write *p*(Δ*R_i_*(*k*)) Similarly, where k goes from 1 to *n*where *n*is the number of states that define *ΔRi*. Similarly *p*(Δ*R_i_*(*k*)),(Δ*R_j_*(*l*)). In order to simplify the notation, we will suppress the state index, and write *p*(Δ*R_i_*(*k*)) as *p*(Δ*R_i_*). Similarly, *p*(Δ*R_i_*(*k*),Δ*R_i_*(*l*)) ≡ *p*(Δ*R_i_*,Δ*R_j_*).

### Time delayed correlation functions

We let *p*(Δ*R_i_*(*t*)),(Δ*R_j_*(*t+τ*)) denote the joint probability of observing the fluctuation *ΔR_i_* at time t and *ΔR_j_* at time *t+τ*. In this simplified notation, *ΔR_i_(t)* represents the value of *ΔR_i_* in state k at time t,
which is identical to Δ*R_i_* (*k*, *t*).Δ*R*_*j*_ (*t+τ*) may be affected by the earlier fluctuations of Δ*R_i_* (*t*). The extent of this effect may be quantified by the time delayed correlation function

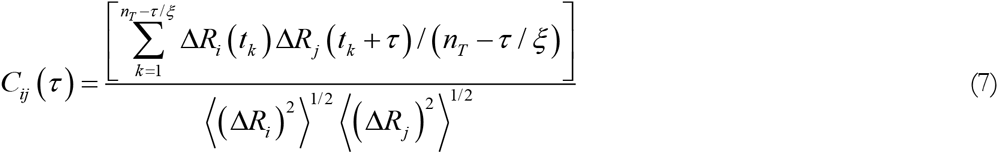

This is a conditional correlation where Δ*R_i_* comes before Δ*R_j_*. In general, 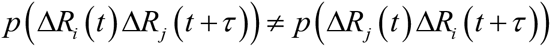. This leads to directionality in the structure, known as causality, and consequently,

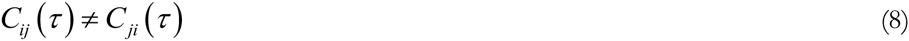

If the fluctuations of residue *i* control the fluctuations of residue *j*, i.e., if residue *j* is driven by*i*, thenthe decay time for *C_ij_*(τ) will be larger than that of *C*_*ji*_(τ).

When τ = 0 time independent Pearson correlation function is obtained as

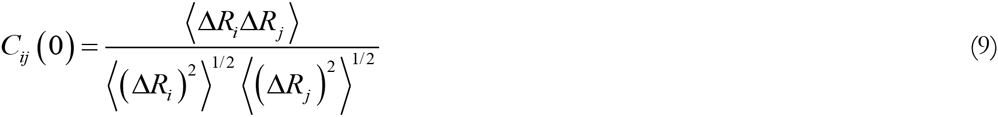

### Entropy

The entropy for a system with pair probabilities is given by

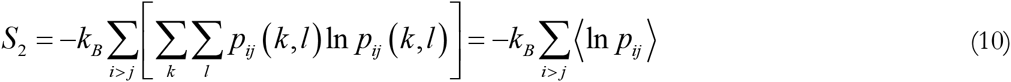

For brevity of presentation, we used the notation 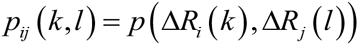 In Equation 10,*S*_2_ signifies the joint entropy for a system with pair probabilities.

We now divide and multiply the entropy expression by the singlet probabilities:

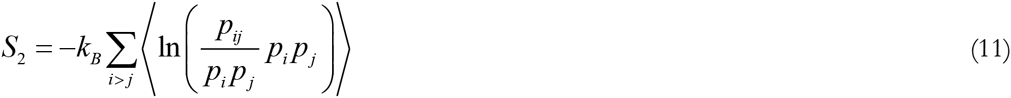

and expand Eq. 11 as

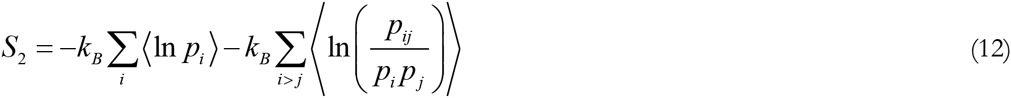

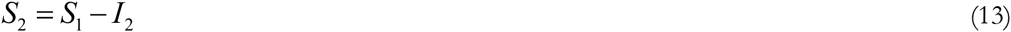

where, 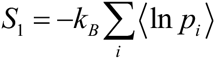 is the singlet entropy and 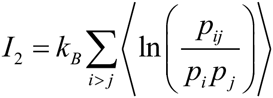 is the mutual information of the system.

Equations 10 to 13 are written for the full system. Mutual information is more meaningful when two subsystems of the protein are of interest. Using statistical mechanics arguments given by Callen [24], each subsystem may be treated as a canonical ensemble that exchanges energy with its surroundings, represented by the cartoon in Fig 12. The surroundings of Subsystem 1 for example is the protein which contains Subsystem 2 also. We may choose the subsystems arbitrarily, an atom, an amino acid, or a secondary structure such as a helix, beta strand, loop or a tail. The subsystem may also be in contact with the surroundings of the protein. Mutual information is zero if the fluctuations of *i* are independent of the fluctuations of *j*. Otherwise, mutual information is always greater than zero. This leads to the conclusion that correlations always decrease the sum of the individual entropies in a system.

**Fig 12.**
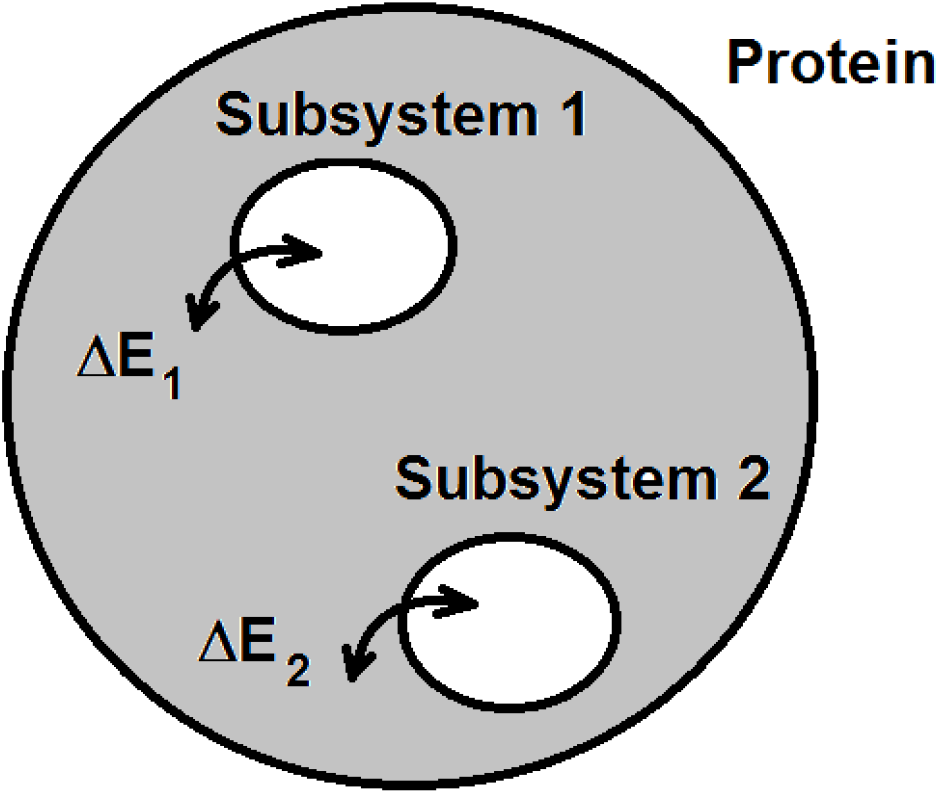
*Energy exchange between protein and its subsystems.*

### Conditional entropy

We consider two trajectories, Δ*R_i_*(*t*) and Δ*R_j_ (t)*. We now consider two events separated in time by τ, with the condition that Δ*R_i_* coming before Δ*R_j_*. The conditional entropy for these two events is defined by

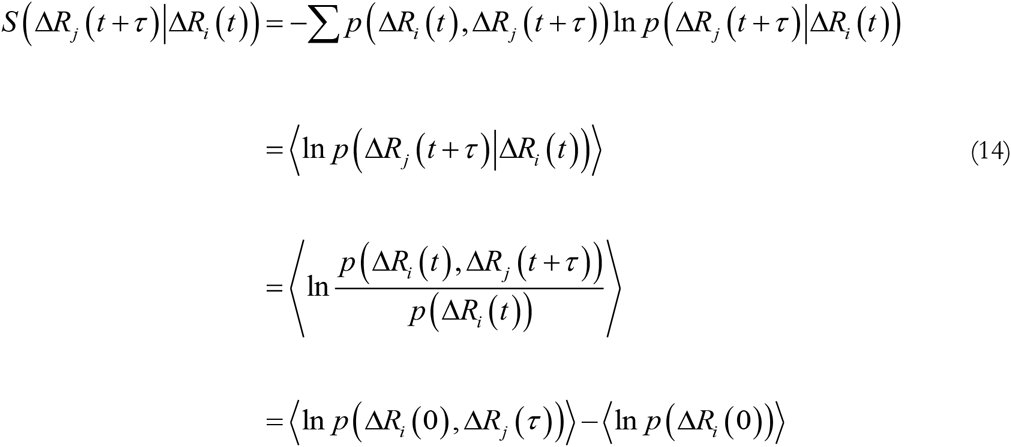

where, the summation is over all states for *i* and *j*, and the condition of stationarity is used in the last equation.

### Transfer entropy

Following Schreiber’s work [22], we write the transfer entropy *T_i→j_*(τ) from trajectory *i* to *j* at time τ as

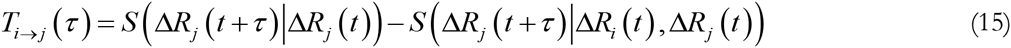

Using the last of Eq. 14, this may be written as

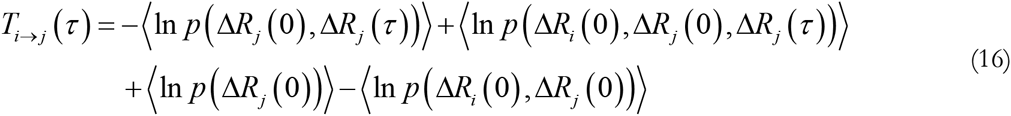

Through the term 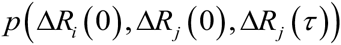, Eq. 16 requires the evaluation of triple probabilities. If trajectories i and j are independent, then *T_i→j_*(τ) = 0 entropy transfer from *i* to *j* will be zero. In general, *T_i→j_*(τ)≠*T_i→i_*(τ) and this will determine the net transfer of entropy from one event to another separated in time by τ . Different values of τ shows how entropy transfer depends on prior interactions. In this study, we will take τ=5 *ns* as the representative correlation time of cross correlations.

### Net entropy transfer from an atom

Equation 16 gives the entropy transferred, *T_i→j_*(τ), from atom i to j. The net entropy transferred from to to all other atoms is obtained by summing over all j as

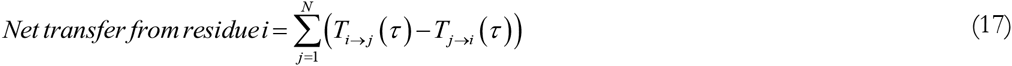

### Molecular Dynamics simulation

All-atom Molecular Dynamics simulations were performed for unbound (PDB ID: 1UBQ) and bound states (PDB ID: 2KTF) of Ubiquitin, using NAMD 2.11 simulation program with CHARMM22 All-Hydrogen Parameter set for Proteins and Lipids. TIP3P water model was used to represent water molecules. Counter ions are placed to neutralize the system. Time step of simulations were 2 fs and periodic boundary conditions were applied in an isothermal-isobaric NPT ensemble with constant temperature of 300 K and constant pressure of 1 bar. Temperature and pressure are controlled by Langevin thermostat and Langevin piston barostat, respectively. System coordinates were saved every 1 ps. 1-4 scaling is applied to van der Waals interactions with a cutoff of 12.0 Å. Energy of the system was minimized and system is heated to 300 K for 50 ps and further subjected to MD production run for 600 ns. Frames in trajectories were aligned to the first frame of the simulation by using VMD 1.9.2 to eliminate all rotational and translational degrees of freedom and the analysis is done with the aligned Cartesian coordinates.

### Entropy Calculations

Amplitude of fluctuations were calculated for each atom from Cartesian coordinates,*R_i_ (t_k_)*, of the thetrajectory and the mean amplitude of fluctuations, 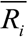 was subtracted from each *R_i_*(*t_k_*) and the ΔR_i¨N_(*t_k_*) matrix was generated with (*t_k_*,*N*) dimensions, where *N* is the number of atoms and *t_k_* is the number of frames selected for calculations. All of the calculations in this study will be based on the alpha carbon of each residue unless otherwise stated. Initial data up to equilibration was excluded from the calculations as equilibration. A binning approach was used to calculate configurational entropy, for individual and pairwise dependent atoms. Calculations were performed using MATLAB R2015b. Histogram function of MATLAB was used to cluster data into 8 bins with specified widths and partitioning of data is adaptive according to the maximum and minimum of data. Calculations were performed using 8 discrete bins. Number of bins were selected according to the Sturges’ rule. The optimum number, nopt, of bins is calculated from the Sturges’ rule according to

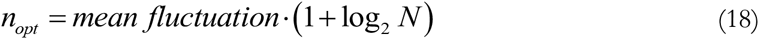

Here, the mean of fluctuations, i.e., the average fluctuation of the *N* alpha carbons is calculated from the trajectories and is in the order of 0.5 Å. For a trajectory of 600,000 time steps, the optimum number of bins is obtained as 8 which is used throughout the calculations. After partitioning the fluctuation of each atom into 8 discrete bins, the probabilities were calculated from the frequency of occurrences and entropy was expressed in individual and pairwise mutual information terms. For comparison with benchmark calculations, the configurational entropy was calculated for all heavy atoms by subtracting pairwise mutual information term from individual entropy term as given by Eq. 12. By using Eqn. 16, transfer entropy from ith atom to j was calculated with a delay value of 5 ns for alpha carbons by subtracting the triple conditional entropies from pairwise conditional entropies. Result of configurational entropy calculations were compared with benchmark data and transfer entropy results were used to study changes in entropy transfer patterns when Ubiquitin forms a complex.

Amount of mutual entropy depends on the distribution of the individual entropies and it is bounded by individual entropy terms.

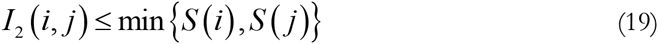

Estimated entropy from a finite sample may be affected by some systematic errors and a correction term is required to get rid of this error [55]. Corrections were applied according to the previous studies [56]

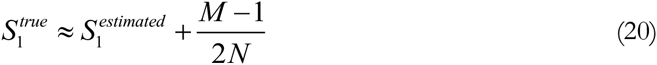

Where *S^estimated^*_1_ is the raw entropy, *M* is the number of histogram bins with non-zero probability. Since mutual entropy is a sum of entropies, this formula can also be used to correct *I*_2_(*i,j*) terms.

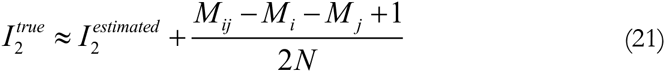

Where *M_ij_*, *M_i_* and *M_j_* represent the numbers of the corresponding histogram bins with non-zero probabilities.

### Benchmark

A benchmark data set which contains one protein-protein complex was used to compare our results with results of MIST(Mutual Information Spanning Trees) method of PARENT [38]. This protein complex is human polymerase iota UBM2 in complex with Ubiquitin, 2KTF.pdb. Unbound state of free Ubiquitin, 1UBQ.pdb, was also simulated. The entropy change upon complex formation was obtained as the difference between configurational entropies of the unbound and bound states.

**Fig 13.**
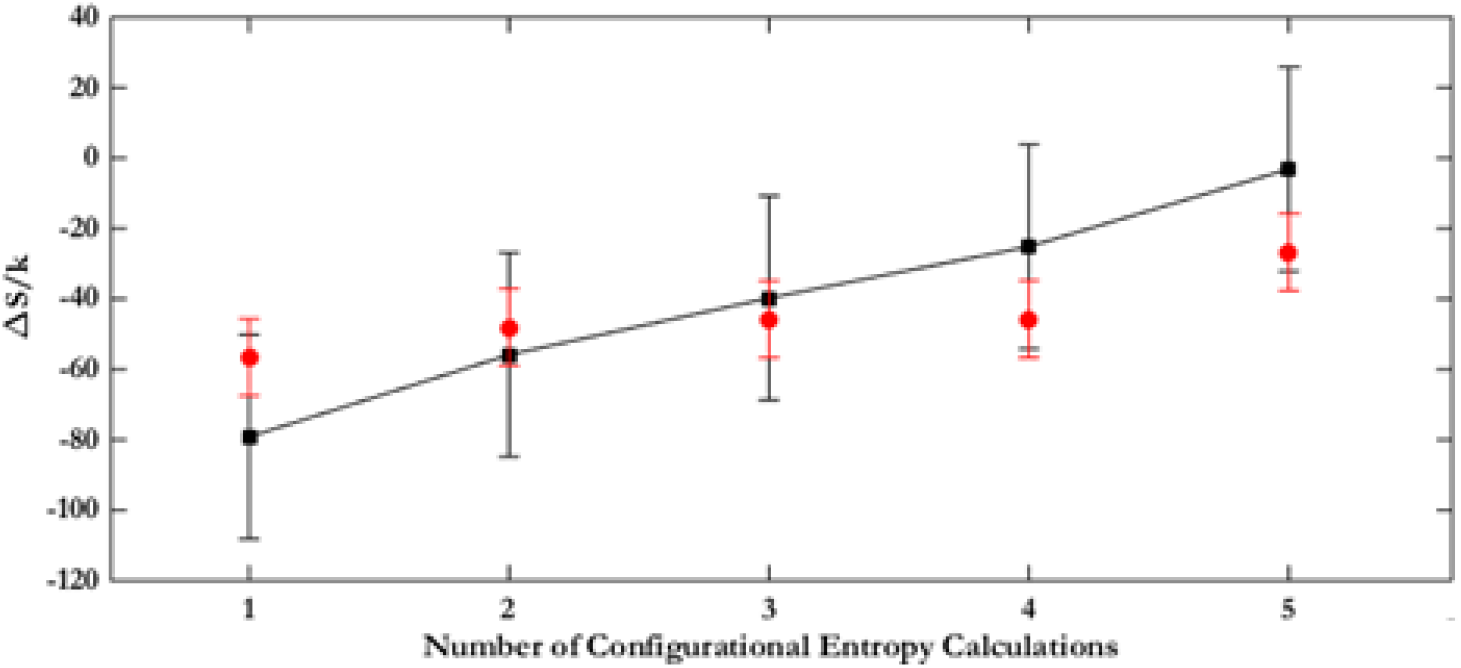
*Comparison of configurational entropy of binding of UBQ and UBM. Black points are from this work, red points are from PARENT. The vertical lines denote the error bars calculated from PARENT.*

The points from this work are in perfect agreement with the results of PARENT.

## Discussion

In the present paper we used the Schreiber’s model of entropy transfer and presented a detailed analysis of allosteric communication in Ubiquitin. Based on the analysis of time delayed events, we showed that information may be transferred between pairs of residues. Ubiquitin is not generally known as an allosteric protein but our work shows that there is significant information transfer between residue pairs in this system. From the entropy transfer point of view, all proteins may exhibit allosteric communication. This observation supports the recent hypothesis by Gunasekaran et al [18] that allostery is indeed an intrinsic property of proteins. Our work shows that the knowledge of time delayed correlations and entropy transfer is needed in order to quantify allosteric communication in proteins. Time delayed events have not been widely used in studies of protein function and allosteric communication. Recently, it was shown that causality introduced by time delayed correlations plays significant role on the allosteric communications in K-Ras [48]. In this respect, time delayed correlation functions may be viewed as a new tool for studying allosteric communication in proteins. A three dimensional map of entropy transfer, as shown in Fig 14, which is a three dimensional version of Fig 3, may be useful for visualizing allosteric communication between pairs of residues more easily. The residue index on the ‘entropy donor’ axis denotes the residues from which entropy goes out, and the residue index on the ‘entropy acceptor’ axis denotes the residues to which entropy goes. For example, residue 76 on the ‘entropy acceptor’ axis shows that entropy goes into residue 76 from almost all other residues. This means that residue 76 extracts entropy from the protein. Similarly, the peak at residue 53 on the ‘entropy donor’ axis and residue 23 on the ‘entropy acceptor’ axis denotes that residue 23 extracts entropy from residue 53.

**Fig 14.**
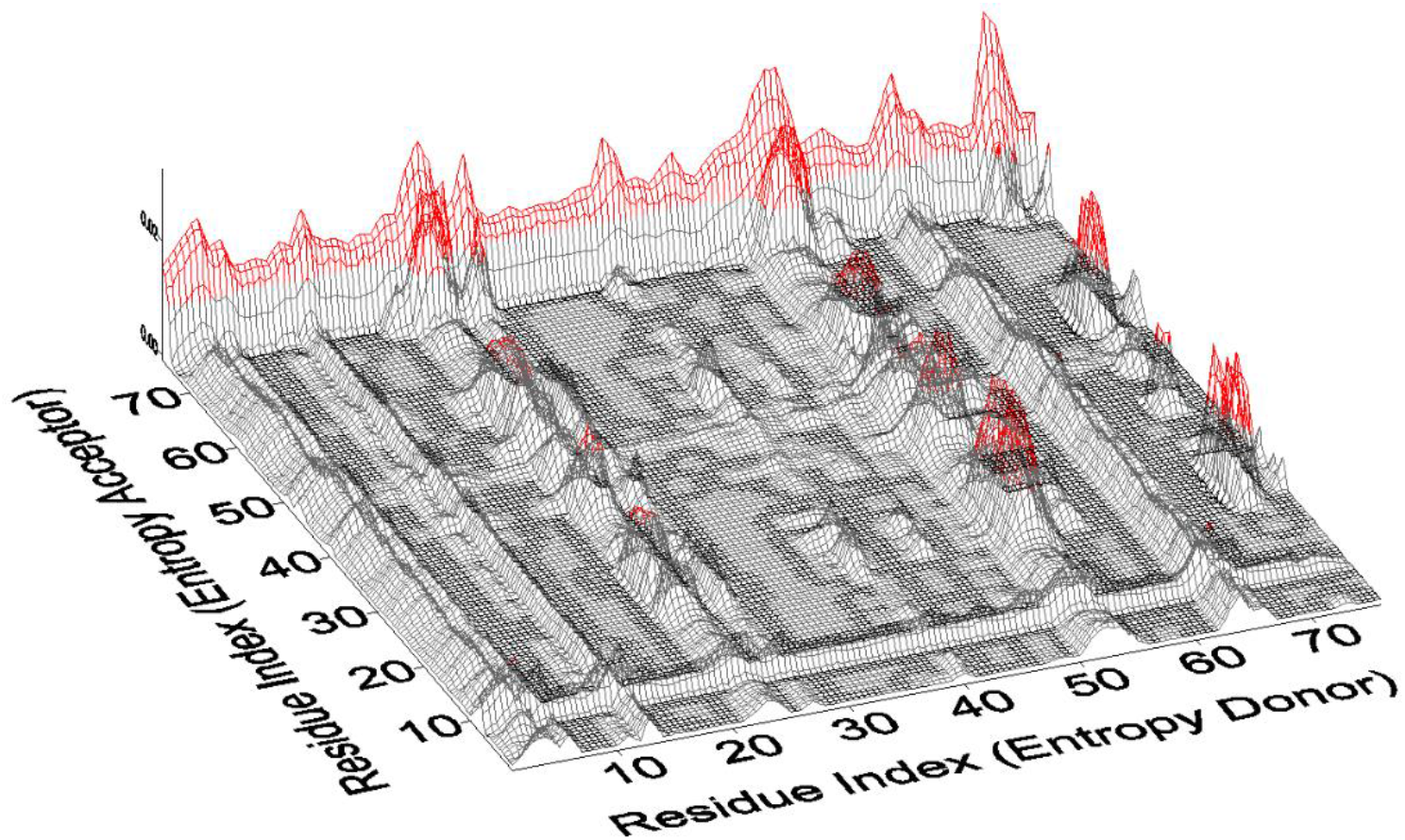
*Three dimensional description of entropy transfer in Ubiquitin. Red regions denote the residues with large contributions to transfer entropy. The figure is a 3-d version of Fig 3.*

Finally, it is worth noting that the present approach which maps the causality, driver-driven relations, and entropy exchange into pairs of residues, as seen in Fig 14, should be of great significance for allosteric drug design because it tells us which residues to manipulate. In this respect, a driver residue is more critical than the driven residue and manipulating the driver will be perturb the existing correlations more efficiently. The effects of mutation on allosteric communication may be quantified by calculating the changes in entropy transfer. As we showed in the UBQ-UBM complex, binding may result in entropy changes in the exposed residues of the complex and change the binding propensities of the complex to other molecules such as another protein, a small molecule ligand or a DNA segment.

